# Social inequity aversion and fairness preference in polar bears

**DOI:** 10.64898/2026.09.07.746566

**Authors:** Kristle Kranenburg, Filipe C. R. Cunha

## Abstract

Decision-making is key to survival, with choices and behaviour typically shaped by evolutionary pressures to enhance fitness and minimize loss. Inequity aversion, the tendency to respond negatively to unequal reward distributions, is therefore unexpected, as acts of fairness may be costly to the individual in the short term. In social, cooperative species, however, fairness may enhance long-term benefits through stable social interactions. Inequity aversion is believed to promote cooperative social structures and has only been demonstrated in certain social species. Yet, solitary species have not been tested, and whether this behaviour depends on social cooperative structures is unknown. Examining solitary species could clarify whether inequity aversion reflects an adaptation to cooperation or a general cognitive capacity linked to social comparison and reward evaluation. We studied inequity aversion in polar bears (*Ursus maritimus*), a largely solitary species, using three complementary, non-invasive cognition tests on seven bears from two zoos in the Netherlands. (1) An impunity experiment assessing effort-based inequity, in which bears performed a standardized action to obtain food rewards. (2) A choice-based experiment testing active selection between equal and unequal reward distributions. (3) A no-task control experiment assessing responses to unequal reward distribution without effort. Under effort-based inequity, bears reduced task performance and increased task latency when a partner received a superior reward. Unequal reward distributions did not affect reward acceptance independent of effort. In the choice-based paradigm, bears preferentially selected equal reward distributions with a partner present. Although generalizability to wild populations is limited, these findings demonstrate behavioural aversion of social inequity in a non-cooperative species. Inequality aversion may therefore not be restricted to social, cooperative animals but instead reflect a broadly distributed cognitive mechanism that emerges under relevant social conditions. These findings challenge current theories and advance our understanding of the cognitive foundations of fairness-related behaviour in animals.

## 1. Introduction

Decision making by humans is not solemnly based on maximizing gain or minimizing loss, but also on fairness (Fehr & Schmidt, 2011). Humans show a natural preference for equal reward distributions. When two or more individuals are unequally paid after performing an identical task, the receiver of the lesser reward demonstrates negative responses (Ostojić & Clayton, 2013; Roth, 2020). The ability to recognize social inequity and a sensitivity towards unfairness may have played a key mechanism in the evolution of human cooperation (Brosnan & De Waal, 2003; Fehr & Fischbacher, 2004; Stevens & Hauser, 2004; Brosnan et al., 2010; Silk & House, 2011). Other animal species have cooperative social structures as well and in 2003, Brosnan and de Waal demonstrated that brown capuchin monkeys (*Cebus apella*) also respond negatively towards unfairness. Under experimental conditions, capuchin monkeys who witnessed a partner receiving a higher valued reward after performing an identical task displayed frustration or refused task re-execution (Brosnan & De Waal, 2003). Particularly when the conspecific was rewarded without performing the task at all (Brosnan & De Waal, 2003).

Social inequity aversion seems to be a trait of socially cooperative species, rather than a unique human trait, and it may stem from a single origin or has evolved convergently across complex cognitive species with a social lifestyle. The evolutionary development and the mechanisms underlying fairness can be considered from studies on group-living species from different phylogenies, including: bonobos, chimpanzees, cleaner fish, crows, dogs, gorillas, macaques, marmosets, keas, orangutans, owl monkeys, rabbits, rats, squirrel monkeys, tamarins, and wolves (Oberliessen & Kalenscher, 2019 – Table 1). The results across and within species vary. For example, dogs demonstrate resentful behaviour towards conspecifics receiving superior rewards (McGetrick & Range, 2018) whereas cleaner fish show no social inequity aversion behaviour (Raihani & McAuliffe, 2012). In rats, social inequity aversion is observed consistently (Hernandez-Lallement et al., 2015; Marquez et al., 2015; Hernandez-Lallement et al., 2016; Oberliessen et al., 2016; Hernandez-Lallement et al., 2018), while for Chimpanzees, some experiments do indicate social inequity aversion (Brosnan et al., 2005; Brosnan et al., 2010; Horner et al., 2011) and others do not (Jensen et al., 2007; Brauer et al., 2006; Brauer et al., 2009; Kaiser et al., 2012). Hence the existence of inequity aversion in non-human animals is still debated (Ritov et al., 2024)

**Table 1.**
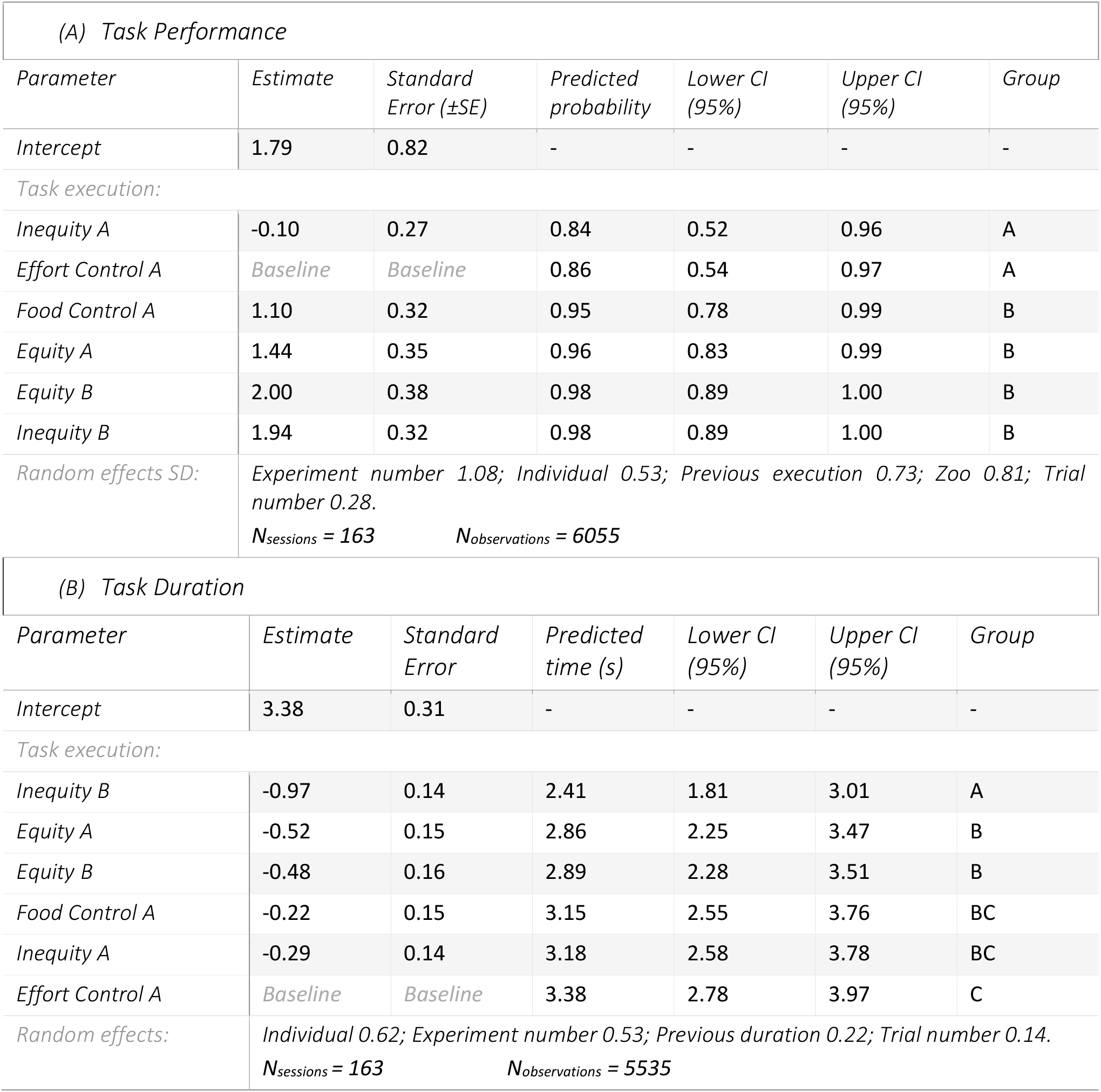
Parameter estimates of the best GLMMs modelling (A) task performance and (B) task duration under impunity experimental conditions. For fixed factors, model estimates, standard error (±SE), predicted values with 95% CI and Tukey post hoc groupings are shown. Random effects are presented as standard deviations (SD).

Multiple social and non-social motives presuppose how an individual reacts to unfairness and may partly underly the heterogeneity in findings (Oberliessen & Kalenscher, 2019; Ritov et al., 2024). For instance, Roma et al. (2006) used a modified set-up from Brosnan and De Waal (2003) to test capuchin monkeys and found that individuals receiving a more valuable reward at the beginning of the experiment but a lesser reward at the end exhibit higher rejection rates, regardless of their conspecific’s reward. Here, rejection more likely represents frustration due to deprivation of the more valuable reward than that it reflects inequity aversion. Various constructs, including frustration, social disappointment, loss aversion, reward expectation, reference-point dependency, and fairness preference, are relevant in understanding social inequity aversion behaviour. These constructs are not mutually exclusive but work in concordance (Oberliessen & Kalenscher, 2019). A task design that balances and counterbalances these theories is essential and to account for them. Oberliessen and Kalenscher (2019) recommend a combination of experiments including impunity and choice-based tasks. Impunity-based tasks involve costly decisions and external reward distributions decided by the observer, while choice-based tasks are cost-free and regard reward distributions based on the subject’s choice. Thus far, only Jensen et al. (2007) and Kaiser (2012) (on chimpanzees) conducted such comprehensive approach.

Current theory suggests that inequity aversion only occurs in socially cooperative animals, however, there is a lack of studies that have examined the trait in solitary species such as snakes, tigers, or bears. Studies on solitary species could support or improve current theory and clarify evolutionary origin, functional role, and underlying mechanisms of inequity-related behavioural responses. In this study, we therefore tested inequity aversion in polar bears (*Ursus maritimus*).

Polar bears are considered solitary species as they tend to live and hunt alone, minimize social interactions, and keep distance from conspecifics (Latour, 1981; Nowak, 1999; Armstrup, 2006; Renner & Kelly, 2006; Davis & Pagano, 2021). Though their solitary lifestyle is interrupted during mating season and the three-year parenting period between mother and cubs (Renner & Kelly, 2006; Davis & Pagano, 2021). Other social interactions are limited to aggregations at highly concentrated food resources, such as whale carcasses and city dumps, or during summering on land (Stirling et al., 1977; Renner & Kelly, 2006; Davis & Pagano, 2021). Given their solitary lifestyle and lack of social cooperative structures, polar bears would not be expected to exhibit social inequity aversion. However, polar bears have a high cognitive functioning, they perform tool-using behaviour, which only occurs in very few non-primates (Deecke, 2012; Stirling et al., 2021), and show marked awareness of conspecific behaviour. By assuming that cognitive capacity and social awareness form foundational components of fairness perception, rather than social structures, polar bears are expected to be able to detect inequity and respond behaviourally to unfairness.

To assess social inequity in polar bears, we conducted a series of complementing cognitive experiments: an impunity-, a choice-based-, and a no-task control paradigm. These paradigms focussed in varying degrees on: (1) task performance; (2) reward acceptance; and (3) self-chosen reward distribution preferences, under different levels of fairness. If polar bears exhibit inequity aversion, we predicted that: (1) unequal reward distributions following task performance would increase task refusal and prolong task duration; (2) under unfair conditions, lower-value rewards would have higher rejection rates; and (3) when given a non-costly choice, individuals would prefer equal reward distributions over unequal ones.

## 2. Subjects and Testing Procedures

We selected nine polar bears (male and female) from long-term captive groups housed in three Duch zoos: Eindhoven zoo (2F, 1M), Ouwehands Dierenpark (3F, 1M), and Aqua Zoo (2M). After completion of the first experiment, we excluded the two bears from Aqua Zoo as one bear had to withdraw from participation. The final study group therefore consisted of seven bears ranging in age from 1.5 to 27 years. In each zoo group, the oldest individual was wild-caught, and the remaining bears were their offspring.

We tested bears in dyads consisting of a focal bear and a partner. We conducted all experiments in the indoor part of the enclosures, positioning dyads opposite or sideways to each other to allow visual, vocal, and olfactorily contact and to ensure reciprocal observation of task performance and reward delivery. Each session for an experimental condition comprised 25 trials per individual. Within each condition, both bears in a dyad completed the experiment twice as focal subject and twice as partner. Thus, for every partner combination, an individual served as focal subject in two sessions and as partner in two corresponding sessions. Consequently, individuals completed six focal sessions per condition in Ouwehands Dierenpark (2 x 3 partners) and four focal sessions per condition in Eindhoven Zoo (2 x 2 partners), resulting in 96, 48 and 48 impunity, choice-based and no-task sessions in Ouwehands, respectively, and 48, 24, and 24 sessions, in Eindhoven zoo, respectively.

After a training period and prior to each of the three experiments, we assessed the eligibility of an individual in a solo pre-test consisting of 25 trials. A bear qualified if achieving a task success rate of > 90% (n ≥ 23 trials). During these pre-tests, we distributed only the lesser reward (herring in Ouwehands Dierenpark; bread in Eindhoven Zoo). Across all zoos, cod oil served as the superior reward. The rewards were removed from the daily diet, if formerly included, and we assured not to cause any type of nutritional deficiency. Zoos maintained their standard feeding schedules (daily in Eindhoven Zoo and every other day in Ouwehands Dierenpark). However, on experimental days, if applicable, food distribution was delayed until after the experiments to create a minimal food deprivation.

We alternated the order of experiments per zoo and randomized the sequence of test conditions and subjects as far as situational constraints allowed. Situational constraints included day-to-day variations in individual health and wellbeing, which were monitored by experienced zookeepers who maintained daily contact with the bears.

We recorded all experimental sessions on video from which we obtained measures of task performance, such as choice, reward acceptance, and latency. Due to noise and camera angle constraints within the enclosures, we disregarded behavioural reactions (e.g. growling, withdrawal, or lethargy). Per zoo we scheduled two to four sessions per experimental day, although this could vary due to practical constraints causing some individuals to be tested more than once a day. Excluding the training phase, we conducted 65 experimental days between May 26th and November 26th, 2021, working in one or both zoos on a given day.

Additionally, as a precaution to prevent redirect aggression, we made sure a permeable barrier would separate a sideways positioned dyad to limit tactile contact to avoid potential injury (Renner & Kelly, 2006). However, we observed no noteworthy aggressive interactions during the study. Furthermore, subjects could leave the experiment at any time and roam freely through the enclosure to maintain welfare, which we monitored throughout the study. Lastly, during the study, two bears developed skin irritations of unknown origin and received veterinary treatment (Apoquel and Prednisolon Mylan). We temporarily excluded one individual from the choice-based experiment to ensure her wellbeing. After recovery, we repeated a limited number of sessions to compensate for missed trials and to maintain data robustness.

## 3. Experimental Outline

### 3.1 Impunity experiment

In the impunity experiment, we examined how how polar bears respond towards unequal reward distribution after performing an energy-costly task. The experiment consisted of four conditions: *equity, inequity, food control*, and *effort control*. The *equity* condition served as a baseline for fair circumstances: both bears performed the task, and both bears received the lesser reward. An unfair situation was created in the *inequity* condition: both bears performed the task, but the partner received the superior reward, while the focal bear received the lesser reward. To distinguish responses to social unfairness from responses to mere visibility of inaccessible (superior) food, we included a *food control* condition in which the partner was absent and the superior reward was delivered to an empty enclosure as if the conspecific was present. Finally, to assess whether partner effort influenced inequity responses, we added an *effort control*. Here, the partner received the superior reward, without performing the task, while the focal bear performed the task and received the lesser reward.

For the task, bears were required to raise on their hind legs with both front paws resting on the contact window of the enclosure. We selected this behaviour in consultation with the zoo staff because it facilitates medical inspections of paws and fur. An experimenter requested the behaviour using a standardized verbal cue and hand signal, alternating between bears. We scored the task as a refusal if the bear did not perform it within 15 seconds. Because none of the bears had previously been trained for this behaviour, we preceded with a training phase before the experimentation period.

### 3.2 Choice-based experiment

To test whether bears actively avert from creating inequity when given the opportunity, we conducted a choice-based experiment. In this paradigm, the focal bear always received the superior reward, while its choice determined whether the partner received an equal (superior) or unequal (lesser) reward.

We presented two visually distinct targets, which differed in both colour (yellow vs. blue) and shape (star vs. circle). Each target was consistently associated with one of the two partner reward outcomes. To control for potential colour or shape biases, we changed the colour–shape combinations and their associated reward meanings across zoos. Therefore, the target linked to the equal or unequal reward differed between institutions.

Bears indicated their choice by touching the target with the snout or paw. The experimenter held the targets against the enclosure’s contact window to ensure accessibility. To control for side biases, we alternated target position and switched targets between hands systematically. The same experimenter conducted sessions across zoos to maintain consistency in presentation and randomization procedures.

To allow subjects to learn the association between each target and its corresponding partner reward, we conducted control trials in the absence of a partner. During these trials, we delivered the associated reward to an empty enclosure as if a bear was present. We used these controls as individual baselines to account for potential intrinsic preferences for specific visual target features.

#### No-task experiment

In the no-task experiment, we examined responses to unequal reward distribution in the absence of effort. No work in the form of a task was required and hence, rewards were distributed freely to a dyad under two alternating conditions: *equity* and *inequity*. In the *equity* condition, both bears received the lesser reward. In the inequity condition, the focal bear received the lesser reward while the partner received the superior reward. This design allowed us to isolate the effect of unequal reward distribution without the confounding influence of task-related effort.

## 4. Ethic Statement

For this study we complied with Dutch government guidelines regarding animal testing. Permission to conduct non-invasive cognition experiments was received from all participating zoos and Wageningen University. None of the experiments were invasive as described by the Dutch law on animal testing (Wet op de dierproeven, 1977), and hence no experimental license was required. Animals were taken care of by experienced animal caretakers, who ensured none of the animals to be harmed by the experimental procedures. All individuals entered the experimental compartments voluntarily and could leave at any given time.

## 5. Analysis

### 5.1 Video analysis

We analysed approximately 51.5 hours of video footage, including ~36.5 hours from the impunity experiments, ~9.0 hours from the choice-based experiments, and ~6.0 hours from the no-task experiments.

From impunity experiment videos we recorded task execution (1 = performed, 0 = refused), reward acceptance (1 = accepted, 0 = rejected), and task duration (in seconds). We defined task execution as successful when the bear completed the task within 15 seconds of the cue, and as refusal when the bear refused or failed to do so in this time limit. We scored reward acceptance as accepted when the subject consumed the reward, and as rejected when the subject pushed the reward away, or spit it out, leaving more than 50% uneaten. We measured task duration from the moment the experimenter initiated the request (marked by hand gesture reaching its highest point) until the subject made contact with the reward.

From the choice-based experiment recordings, we noted choice outcome: fair distribution or unfair distribution of rewards. From the no-task experiment videos, we recorded whether the reward was accepted or refused.

The video analysis was conducted semi-blind to minimize observer bias. To achieve this, we randomized the video order, anonymized subjects, and effaced both experimental condition and subject role (focal or partner). After completing all behavioural scoring from the videos, we merged the coded data with the corresponding metadata.

### 5.2 Data analysis

We conducted all statistical analyses in R (v4.0.5; R Core Team, 2021). Chi-square tests were used to assess the data and followed up by general linear mixed models (GLMM) for more in-depth analysis, if applicable, to assess the effects of experimental conditions on task performance: reward acceptance, task duration, and choice behaviour. We used the *glmer* function for binomial data and *lmer* for continuous data. We compared multiple models using Akaike’s Information Criterion (AIC) and selected the model with the lowest AIC, unless competing models differed by less than 2 AIC units, in which case both were considered (Burnham et al., 2011; Richards, 2005).

For the impunity experiment, we analysed three dependent variables separately: task execution (binomial: 1 = performed, 0 = refused); reward acceptance (binomial: 1 = accepted, 0 = rejected); and task duration (continuous, seconds; only completed tasks were included). For the choice-based experiment, the dependent variable was choice (binomial: 1 = fair, 0 = unfair). The independent variable in both experiments was the treatment condition. For the impunity experiment, we also included the subject’s role (focal vs. partner) as a fixed factor.

We included relevant random effects to account for repeated measures and non-independence: individual, dyad, trial number, experiment number, previous task performance or choice, and zoo. Target colour and shape were initially considered in exploratory analyses but were excluded from the main choice-based models due to collinearity with zoo and strong individual preferences.

Only sessions in which each animal performed at least 15 trials and where partners completed ≥ 90% of tasks were included. We did this to exclude sessions where external influences caused the focal animal to perform less due to, e.g., weather condition such as a thunderstorm, rather than the experimental condition. Model performance and residuals were visually inspected, and post-hoc comparisons were conducted using Tukey HSD tests when appropriate. Predicted probabilities were calculated for interpretation of binomial outcomes.

## 6. Results

### 6.1 Impunity results

We first examined how polar bears responded to unequal reward distribution after completing energy-intensive tasks. Task performance and duration were negatively affected under unfair conditions (Chi-square: X^2^ = 507.23, df = 6, p < 0.001; X^2^ = 6956, df = 27, p < 0.001). Reward acceptance was unaffected, with only two rejections across all trials.

Task performance was best modelled by a GLMM including treatment condition and role (focal vs. partner) as fixed effects, and individual, trial, experiment, previous task, and zoo as random effects. The model showed that bears were less likely to perform the task under inequity and effort control conditions (predicted probability: 0.84–0.86) compared with equity or food control conditions (0.95–0.98; Tukey HSD, p < 0.05) (Table 1A; Figure 1A).

**Figure 1.**
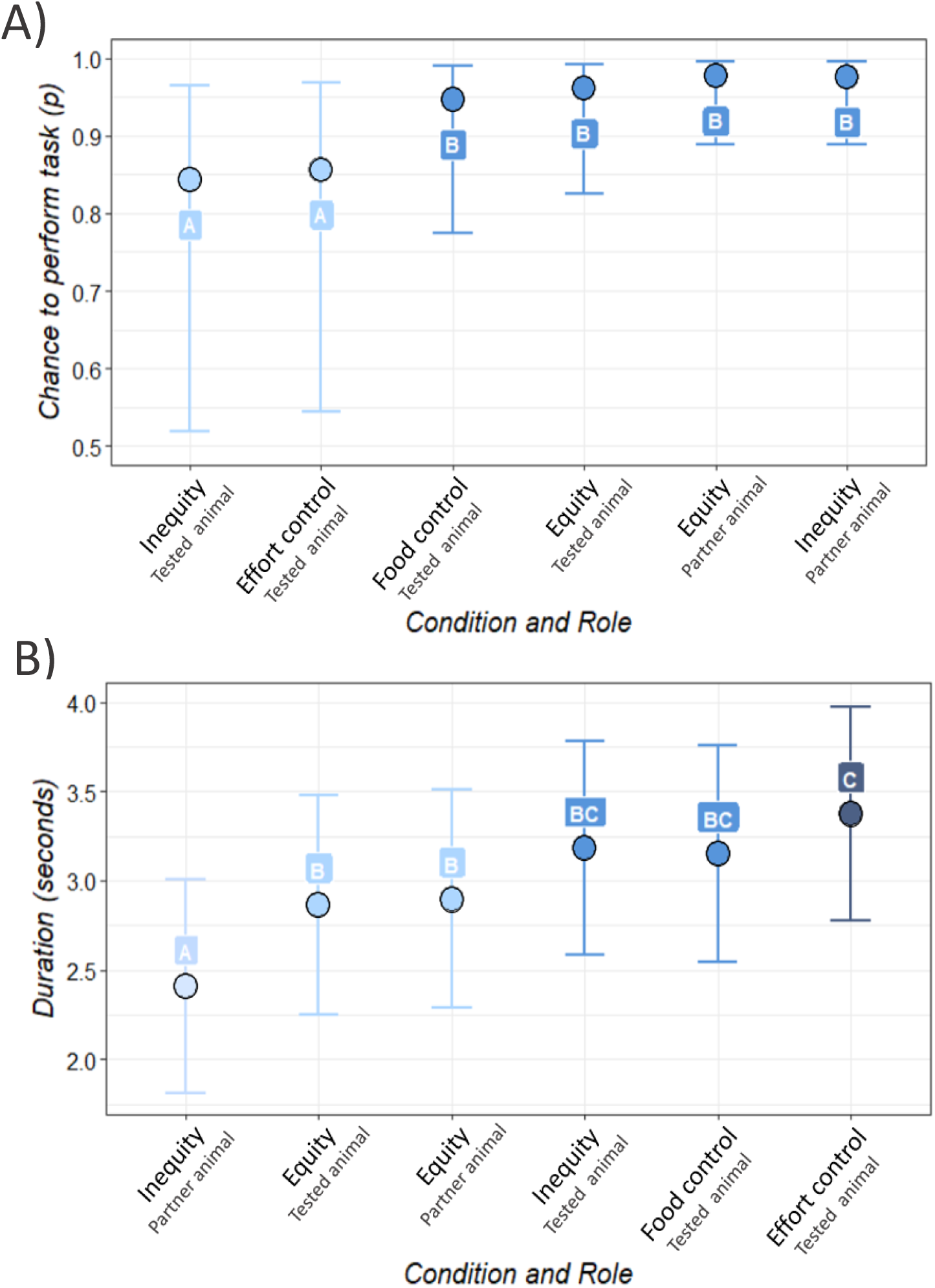
Predicted task execution probability (A) and task duration in seconds (B) of polar bears under impunity experimental conditions, separated by subject role (focal vs. partner), based on GLMM analyses. Points represent model-predicted values and error bars indicate 95% confidence intervals. Different letters denote statistically significant differences according to Tukey HSD post-hoc comparisons. For task execution (A), two significantly different groups were found: *group A* (light blue) and *group B* (dark blue). For task duration (B), three statistically distinct groups were identified (A–C; light blue to dark blue); categories labelled “BC” do not differ from groups B or C.

The task duration, if performed, was similarly influenced by condition and role (GLMM). Bears receiving the superior reward in inequity conditions performed tasks fastest (mean ± SE: 2.41 ± 0.14 s), while bears in the effort control condition performed slowest (3.38 ± 0.31 s). Tasks under equity conditions had intermediate durations (2.86–2.89 s), and bears under inequity and food control conditions fell between the medium and slow groups (3.18–3.38 s) (Table 1B; Figure 1B).

### 6.2 Choice-based results

In the choice-based experiment, results indicated focal bears to prefer equitable outcomes in the presence of a partner (Chi-square: X^2^ = 49.843, df = 1, p < 0.0001; predicted probability: 0.61) than in control trials without a partner (predicted probability: 0.48; GLMM, treatment condition as fixed factor, previous choice and zoo as random effects; p < 0.05) (Table 2, Figure 2.).

**Table 2.** Parameter estimates of the best GLMM modelling choice under choice-based experimental conditions. For fixed factors, model estimates, standard error (±SE), predicted probabilities to choose fair reward distributions with 95% CI and Tukey post hoc groupings are shown. Random effects are presented as standard deviations (SD).

| Parameter | Estimate | Standard Error | Predicted probability | Lower CI (95%) | Upper CI (95%) | Group |
| --- | --- | --- | --- | --- | --- | --- |
| Intercept | -0.07 | 0.25 | - | - | - | - |
| Treatment: |  |  |  |  |  |  |
| Control | Baseline | Baseline | 0.48 | 0.36 | 0.60 | A |
| Choice | 0.53 | 0.08 | 0.61 | 0.49 | 0.72 | B |
| Random effects: | Standard Deviation |  |  |  |  |  |
| Zoo | 0.11 |  |  |  |  |  |
| Previous choice | 3.01·10<sup>-4</sup> |  |  |  |  |  |
| N<sub>sessions</sub> = 122 N<sub>observations</sub> = 2820 |  |  |  |  |  |  |

**Figure 2.**
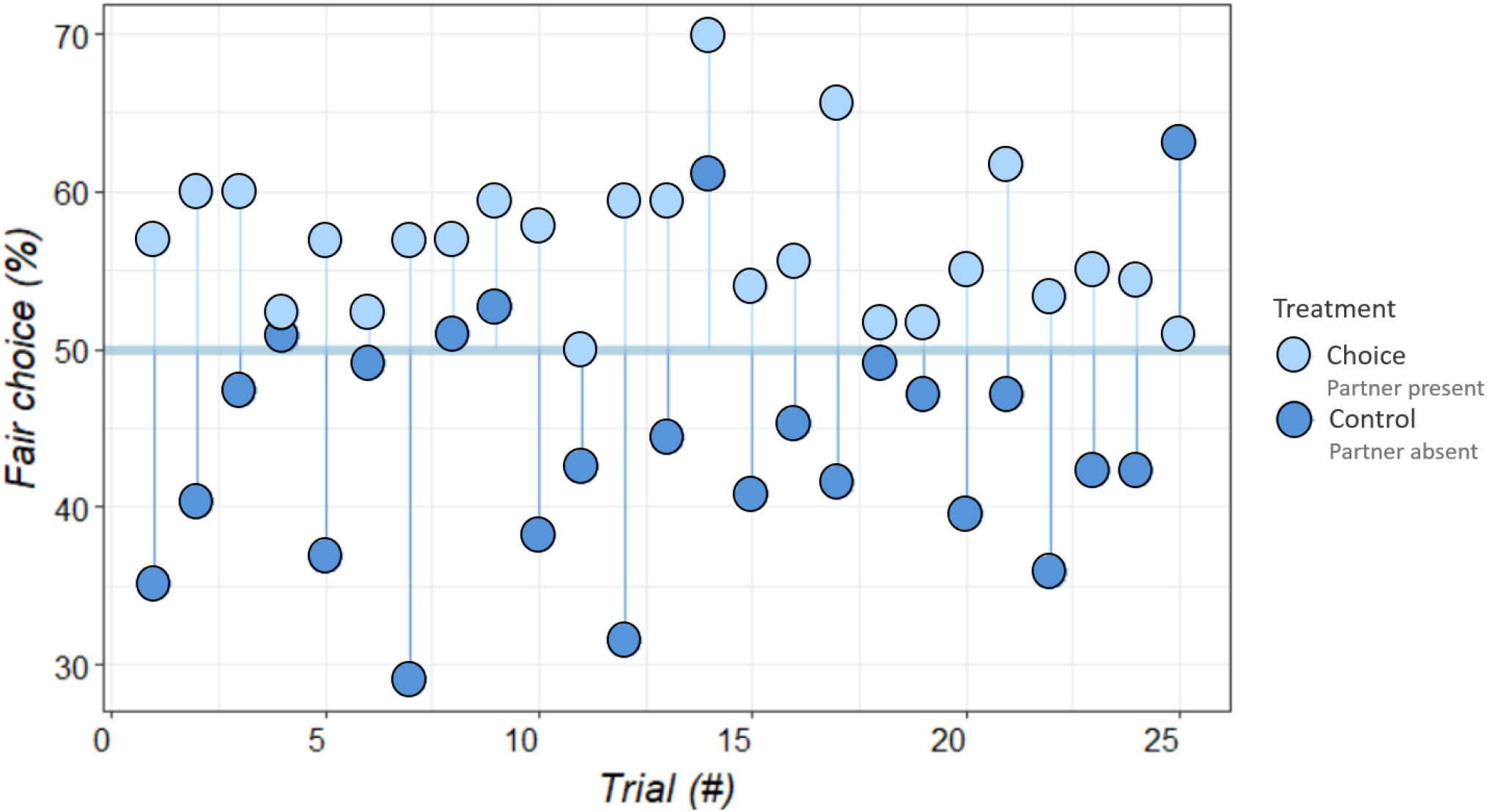
Percentage of fair choices under control (dark blue) and choice (light blue) conditions across trials in the choice/based experimental sessions.

### 6.3 No-task results

When no task was required, bears accepted rewards almost universally, regardless of equity or inequity (equity: 99.2%; inequity: 98.8%; N_sessions_ = 63 N_observations_ = 3176). No further statistical tests were conducted due to the near-uniform acceptance.

## 7 Discussion

Social inequity aversion has been documented in several species, yet the mechanisms and function behind this trait remain debated (Oberliessen & Kalenscher, 2019; Ritov et al., 2024). By convention, inequity aversion is associated with only socially cooperative species. Our findings challenge this assumption. Across complementary experimental set-ups, polar bears, a predominantly solitary species, demonstrated behavioural responses consistent with social inequity aversion.

In the impunity paradigm, bears showed increased task refusal and prolonged task duration under unequal reward distributions. Notably, task refusals increased only when a partner received the superior reward, not when the reward was delivered to an empty enclosure. This distinction indicates that the bears responded to social unfairness rather than to mere inaccessibility of a better reward. Furthermore, task duration further scaled with the level of unfairness, particularly when the partner received a superior reward without exerting effort. These patterns align with findings in other species, where individuals delay responses, reduce effort, or refuse participation under unequal reward conditions (e.g., Takimoto & Fujita, 2011; Hernandez-Lallement et al., 2016).

The choice-based experiment strengthens this interpretation. When given control over reward distribution, bears preferentially selected equitable outcomes in the presence of a partner. This result suggests that polar bears not only react to unfairness but can also proactively adjust their decisions to promote fairness. In contrast, when no task was required, unequal reward distribution did not affect reward acceptance. Together, these findings indicate that effort plays a role in eliciting inequity responses, consistent with the idea that inequity aversion is closely related to a cost–benefit evaluation rather than simple reward comparison.

The results show a behavioural pattern of disfavour towards inequity under energy-related costs and active decision-making. These patterns resemble results reported in several other taxa, including capuchin monkeys (Brosnan & de Waal, 2003; van Wolkenten et al., 2007; Takimoto & Fujita, 2011), wolves under impunity conditions (Essler et al., 2017), crows (Wascher & Bugnyar, 2013), and rats in choice-based tests (Hernandez-Lallement et al., 2016) and great apes, including chimpanzees, in no-task experiments (Bräuer et al., 2006). However, the literature contains conflicting findings both across and within species (Oberliessen et al., 2019), e.g. chimpanzees, under both impunity and choice-based conditions, report no clear evidence of inequity aversion (Jensen et al., 2007).

A recent meta-analysis concluded that non-human animals do not show inequity aversion in accept–reject paradigms (Ritov et al., 2024). Notably, that analysis focused exclusively on binary refusal tasks and did not include effort-based or active choice paradigms. Because inequity sensitivity may manifest through changes in effort, latency, or prosocial choice, or even physical stresses (e.g. heart and age; Heidary et al., 2012) rather than simple reward rejection, restricting analyses to accept–reject outcomes may underestimate its expression. Our own results illustrate this point: although reward rejection was virtually absent in the impunity paradigm, inequity effects emerged clearly in task refusal and latency. Conclusions about the presence or absence of inequity aversion depends considerably on the behavioural measures and paradigms employed (Oberliessen et al., 2019).

Furthermore, our findings suggest that inequity aversion is not confined to cooperative, in-group living species, as previously assumed. The presence of inequity aversion in largely solitary species raises evolutionary questions. One possibility is that the trait evolved from a single origin and may be expressed as an atavism in non-cooperative species. Inequity aversion may emerge flexibly in response to social exposure, even in species that do not rely on long-term cooperation. Under this view, the trait may reflect a general socio-cognitive capacity rather than an adaptation exclusive to cooperative breeding or group living. Alternatively, the expression of the trait might rely on certain cognitive prerequisites, such as reward comparison, social awareness and cost evaluation. This could also mean the trait can be learnt in social (non-natural) aggregations.

Several limitations of this study do warrant consideration. The study involved a limited number of individuals, housed in non-natural social structures. Social exposure in captive environments differs from natural conditions and could influence behavioural responses to inequity, for example by magnifying their reactions. Generalizability to wildlife is therefore limited. However, previous work indicates that polar bears in captivity retain solitary behavioural patterns comparable to those observed in the wild (Latour, 1981; Renner & Kelly, 2006), suggesting that the responses reported here are unlikely to be artefacts of captivity alone. The consistency of responses across institutions further strengthens the robustness of the observed patterns. Additionally, although adult females were overrepresented due to sample constraints, we did not observe qualitatively different responses among infant males. Nonetheless, future studies with more balanced sex ratios and larger samples would allow stronger inference.

Future research should also incorporate physiological measures to assess internal responses to inequity. Behavioural indicators such as refusal or latency capture overt reactions but may underestimate subtle affective responses. For example, Heidary et al., (2012) found that inequal food distributions in rabbits resulted to negative responses in heart quality and aging. Measures such as heart rate, temperature, or stress-related biomarkers could reveal inequity-related responses even in the absence of observable behavioural change. Integrating physiological and behavioural approaches may help explain inter- and intra-species variability and reduce the risk of false negatives.

Understanding inequity sensitivity has broader implications. If solitary species can detect and respond to unfairness, social lifestyle alone cannot explain the distribution of this trait. Instead, inequity aversion may reflect a more general feature of comparative cognition linked to reward evaluation and social monitoring. These findings contribute to ongoing debates about the evolutionary origins of fairness and cooperation and suggest that the foundations of such behaviour may be more widespread, and more flexible, than traditionally assumed. Moreover, recognizing inequity sensitivity in non-humans, especially non-social species deepens, our understanding of animal cognition and emotional complexity, challenges simplified views of animal intelligence, and informs ethical considerations in animal management and policy.

## 8 Conclusion

Responding to fairness can impose fitness costs, and inequity aversion has therefore been assumed to occur primarily in socially cooperative species. Across an array of experimental paradigms, designed to balance social and non-social constructs that underlie inequity responses, we demonstrate that polar bears, a largely solitary and non-cooperative species, recognize and respond to social inequity.

Polar bears reduced task performance and delayed task execution when a partner received a superior reward, particularly when the partner expended no equivalent effort. Yet we found no response to unequal reward distributions in the absence of effort. When given an active choice, polar bears preferentially selected equitable outcomes. Together, these findings indicate that effort and social comparison shape inequity responses.

Although our study was conducted in captivity, limiting generalization to wild populations, the behavioural patterns align with findings from other taxa and challenge the view that inequity aversion is restricted to stable cooperative societies. Instead, inequity sensitivity may reflect a more broadly distributed cognitive capacity that emerges under relevant social conditions.

Future research should integrate physiological measures alongside behavioural data to better capture affective, or delayed responses to unfairness and clarify variation within and across species. By expanding our understanding of inequity aversion, we refine our understanding of the evolutionary foundations of fairness and broaden our understanding of the cognitive and emotional capacities of animals

## Acknowledgements

Our sincere thanks to dr.ir. Bonne Beerda for his supervision, insights, and continued support throughout this project. We are grateful for the collaboration with Ouwehand Dierentuin, Eindhoven Zoo (formerly Dierenrijk), the institutional supervisors and zoological directors who facilitated access, and enabled this research to take place. Our appreciation especially goes to the dedicated animal caretakers for their essential practical assistance, expertise, and commitment during data collection. Finally, we would like to acknowledge the polar bears whose participation made this research possible and who have taught us a lot.

